# *PdeMIXTA04* triggers epidermal cells of placenta to differentiate into poplar catkins through forming MBW complexes with *PdeMYC* and *PdeWD40*

**DOI:** 10.1101/2022.11.01.514779

**Authors:** Fangwei Zhou, Huaitong Wu, Yingnan Chen, Gerald A. Tuskan, Tongming Yin

**Author notes:** Authors contribute equally to this work (Fangwei Zhou, Huaitong Wu). Corresponding author: Tongming Yin, Key Lab of Tree Genetics and Biotechnology of Educational Department of China, Nanjing Forestry University, 159 Longpan Road, Nanjing, 210037, China.

## Abstract

Differentiation of plant epidermal cells is a keen research topic in plant biology. Our study on *Populus deltoides* revealed that epidermal cells of the female flower placenta protruded to form catkin fibers immediately after pollination. We discovered that *PdeMIXTA04* was explicitly expressed in placenta of female poplar flowers. Heterologous expression of *PdeMIXTA04* in *Arabidopsis* demonstrated that it significantly promoted the leaf epidermal cells to differentiate into trichomes. Compared with the wild type, significant increases in trichome density and trichome branches were observed on leaves of all *35S:PdeMIXTA04* transgenic lines. Furthermore, transformation of *PdeMIXTA04* in the trichomeless *Arabidopsis* mutant *(gl1)* restored trichome development to that of the wild type. GUS expression in poplar, driven by the promoter of *PdeMIXTA04*, also confirmed trichome-specific expression. We then screened a yeast library with *PdeMIXTA04* and captured two interacting genes, *PdeMYC* and *PdeWD40*. Interactions between these two proteins were verified by yeast two-hybrid (Y2H), biomolecular fluorescence complementation (BiFC), dual-luciferase (dual-LUC), and pull-down assays, indicating that PdeMIXTA04 functions through the MYB-BHLH-WD40 (MBW) ternary complex. Our work presents evidence of *PdeMIXTA04* as a candidate gene for editing to resolve catkins associated pollution and provides distinctive understanding of the molecular mechanism triggering differentiation of plant epidermal cells.

## Introduction

The genus *Populus* comprises 30-35 species (Naiman *et al*., 2005) that are widely distribute in the temperate and cold temperate regions (latitude 22-70°N) in northern hemisphere (Müller *et al*., 2013). Poplars are important commercial woody plants due to their rapid growth, robust adaptability, and use in a wide variety of end products (Polle *et al*., 2010; Sannigrahi *et al*., 2010; Polle *et al*., 2013; Doty *et al*., 2009). *Populus* species are dioecious plants, bearing male or female inflorescences on trees of alternate sex. The female inflorescence are arranged in catkins, which assist seed dispersal in middle spring through early summer (Cronk *et al*., 2015) and which can cause severe airborne pollution, with many people allergic to the fluffy fibers attached to the seed coats (Xu, 2021). In addition, catkin formation and maturation requires substantial energy input, consuming considerable amounts of photosynthate every year and leading to tradeoffs in vegetative and reproductive growth (Morag, 2018).

Female catkins in poplar are comprised of flocculent fiber cells 9.5 to 15.9 mm in length and 8 to 11 μm in diameter (Chen *et al*., 2010). When the fiber cells of catkins elongate, the nucleus expand and nuclear replication occurs, but the cells do not divide (Ye *et al*., 2014), replication cycles bypass cytokinesis during mitosis to maintain the single-cell structure of catkin fibers (Li *et al*., 2018). Poplar flower and catkin formation are processes that are spatial-temporally regulated. Flower and catkin maturation involves cell wall biosynthesis, and the MYB, ZF-HD, MYKC, GRF, ERF, C3H, C2H2, bHLH, FAR1, and YABBY gene families are thought to play important roles in catkin formation (Ye *et al*., 2014). However, the molecular mechanism triggering poplar catkin development remains unclear.

The *MYB* genes in plants are known to play essential roles in stress response, root hair density, light and hormone signaling, trichome formation, flower and seed development, as well as in secondary metabolism (Stracke *et al*., 2007; Matias-Hernandez *et al*., 2017). In the MYB gene family, the R2R3 MYB is the largest subfamily and includes 22 subgroups based on sequences variation in their C-termini (Dubos *et al*., 2010). Multiple lines of evidence have confirmed that the *MIXTA* genes, which belong to the ninth subgroup of the R2R3 MYB subfamily, play a significant role in regulating epidermal cells differentiation (*e.g*., Lau *et al*., 2015; Wu *et al*., 2018; Zhao *et al*., 2020). As an example, *AmMIXTA1*, the first *MIXTA* gene, discovered in an *Antirrhinum majus* mutant, is the key regulator triggering petal epidermal cells differentiation into conical cells (Zhou *et al*., 2021; Brockington *et al*., 2013; Ambawat *et al*., 2013; Lau *et al*., 2015; Du *et al*., 2012; Jaffe *et al*., 2007). In *Artemisia annua, AaMIXTA1*, a homolog of *AmMIXTA1*, overexpression results in a significant increase in glandular secretory trichomes (GSTs) (Shi *et al*., 2018). In *Gossypium hirsutum*, down-regulated lines of a *MIXTA-like* gene (*GhMML*), *GhMML4*, results in a reduction in fibers production (Wu *et al*., 2018). All *MIXTA* genes possess the characteristic domain of R2R3 MYB and a conserved motif of AQWESAR**AE*RL*RES (Paterson *et al*., 2012). In addition to the above, the *MIXTA* genes are known to regulate differentiation of epidermal cells in a variety plant tissues, including flowers (Baumann *et al*., 2007; Ambawat *et al*., 2013; Lau *et al*., 2015; Du *et al*., 2012; Jaffe *et al*., 2007), leaves (Jakoby *et al*., 2008; Shi *et al*., 2018), fruit (Zhao *et al*., 2020; Lashbrooke *et al*., 2015), and ovules (Walford *et al*., 2011; Wu *et al*., 2018).

It has been proposed that female flowers on poplar catkins are generated through a similar molecular mechanism as found in cotton fibers. The molecular mechanism triggering development of cotton fibers has been well clarified (Wu *et al*., 2018). In cotton, fibers differentiate from the ovular epidermal cells and are regulated by *GhMML3* and *GhMML4* (Walford *et al*., 2011; Wu *et al*., 2018). However, poplar flowers originate from different plant tissue than that observed in cotton fibers. Therefore, elucidating the genetic mechanism underlying female flower development in poplar catkin may expand our knowledge on the molecular biology of plant epidermal cells differentiation. In this study, we characterize the developmental biology of female catkin formation using a series of experiments to uncover the key gene underlying flower development in poplar. Finally, we elucidated how this gene functions by analyzing a series of proteins interaction.

## Materials and methods

### Plant materials

The carpel of female flowers were sampled at 1 day before pollination (−1), 1, 3, 5, 8, and 15 days after pollination (DAP) from a 20-year-old female *Populus deltoides*. The collected samples were used to characterize the molecular development during flower and trichome development (see below). Leaf buds, young leaves, and bark tissues, used for comparative and control assays, were sampled from clonal ramets of the same tree.

### Observation of poplar catkin development

The details of flower development were observed via scanning electron microscopy (SEM -- a FEI Quanta 200 Scanning Electron Microscope (FEI Co. Ltd, Hillsboro, USA) using a 2-kV accelerating voltage). Carpels at progressive development stages were washed with distilled water, fixed, washed again, and dehydrated, then dissected through the middle axis. The dried and dissected carpels were coated with a thin layer of gold before SEM examination following Singh *et al*. (2015).

### Cloning and analyzing the *MIXTA* genes in *P. deltoides*

The consensus sequences of R2R3 MYB DNA-binding domain and the AQWESAR**AE*RL*RES motif (Albert *et al*., 2011) were used to BLAST against the *P. deltoides* sequences (https://www.ncbi.nlm.nih.gov/genome/?term=Populus+deltoides) to identify the *MIXTA* genes in poplar genome (*PdeMIXTA*). With cDNA templates, we amplified the identified genes. Primers used for genes cloning are listed in **Table S1**. The PCR products were then purified, and the purified products were ligated into Blunt Vector and sequenced by a M13 primer.

Clustal-W (Katoh *et al*., 2013) and GeneDoc (Novo *et al*., 2009) were used to align multiple MIXTA amino acid sequences. In this analysis, manual curation was performed based on positions of the matched amino acids in the R2R3 MYB domain of the PdeMIXTA proteins. To build the phylogenetic tree, we included the *MIXTA* genes from *P. deltoides* (Pde), *A. thaliana* (At)*, Mimulus guttatus* (Mg), *A. majus* (Am), *Solanum lycopersicum* (Sl), *Thalictrum thalictroides* (Tt), *Zea mays* (Zm)*, Oryza sativa* (Os), *Gossypium herbaceum* (Gh)*, Salix purpurea* (Sp), *Eucalyptus grandis* (Eg), and *N. tabacum* (Nt). Phylogenetic tree was constructed with Neighbor-joining (NJ) method in MEGA X64 software (Kumar *et al*., 2018) under Poisson correction and paired gap deletion mode, with 1000 bootstrap replications (Wang *et al*., 2022). Distribution of *PdeMIXTA* genes in *P. deltoides* genome was plotted by using the MapChart software (Zhang *et al*., 2015). Genes with physical distance <100 kb were termed as tandem duplicates (Chai *et al*., 2014).

### Comparative expression of the *PdeMIXTA* genes

Total RNAs used for cDNA synthesis were extracted from the carpel samples collected at −1, 1, 3, 5, 8, and 15 DAP. With the synthesized cDNA, qRT-PCR was performed on the A7500 Fast Real-Time PCR System (Applied Biosystems Co. Ltd, New York, USA) with the gene-specific primers (**Table S1**). The *UBQ* (Potri.014G115100) gene was used as the reference standard based on Wang *et al*. (2015). Gene expression was based on three experimental replicates.

### Heterologous expression of the carpel-specific *PdeMIXTA* genes in *Arabidopsis*

Heterologous transformation of *PdeMIXTA04* was carried out with wild-type of *A. thaliana* Columbia ecotype (Col-0). The CDS sequences of the carpel-specific *PdeMIXTA* genes were recombined into the binary vector pCAMBIA1301. Then pCAMBIA1301-*PdeMIXTA* plasmids were transformed into *Agrobacterium rhizogenes* strain GV3101. Mediated by GV3101, the recombined plasmids were delivered into *Arabidopsis* using the floral dip method (Bent, 2006). The transgenic lines were screened and selected on a 1/2 MS media containing 30μg/mL hygromycin. The positive lines were further confirmed by PCR amplification of the transformed genes. Expression levels of transformed genes were evaluated by RT-PCR.

To examine the phenotypic changes, we produced seedlings with T3 seeds of the transgenic lines and seeds of the wild-type. The first true leaves on 10-day-old seedlings were harvested and a 3.0 mm leaf disc in diameter was sampled from each leaf. Trichomes on each leaf disc were counted using an Olympus SZX10 microscope (Olympus Co. Ltd, Tokyo, Japan) at a 10 magnification. For each transformed gene, we analyzed five transgenic lines and each line included 10 plant duplicates. The trichome numbers was reported as mean ± SD (standard deviation). In addition, changes in morphology of the trichomes were recorded using a FEI Quanta 200 Scanning Electron Microscope (FEI Co. Ltd, Hillsboro, USA).

### Confirmation of the tissue-specific expression of *PdeMIXTA04*

To confirm *PdeMIXTA04* expressed in epidermal cells of poplar placentas, we conducted *in situ* hybridization as follows: antisense and sense probes of 34 bp were designed according to the specific sequence of the 3′ end of *PdeMIXTA04* (**Table S1**). Then, probes were digoxigenin-(DIG) labeled by using the DIG RNA Labeling Kit (Roche Co. Ltd, Basel, Switzerland). The female flower buds collected at 1 DAP were infiltrated with the FAA fix solution (90 ml of 50% (v/v) alcohol + 5 ml of glacial acetic acid + 5 ml of formalin) for 12 h, dehydrated in 99.5% (v/v) alcohol, then embedded in paraffin. Subsequently, the processed flower buds were cut into 50-μm-thick slices using a RM2016 pathological microtome. The *in situ* hybridization was carried out based on Wang *et al*. (2021).

To confirm the promoter of *PdeMIXTA04*-mediated tissue-specific expression, we cloned the promoter of *PdeMIXTA04* (GenBank accession No. OP493866), then ligated it into the pCAMBIA-1301 vector to build a GUS reporter construct (**Fig. S1**). Mediated *A. tumefaciens* GV3101, the established construct was transformed into *Arabidopsis* using the floral dip method (Clough *et al*., 1998). GUS activity in transgenic *Arabidopsis* was examined by histochemical staining as described by Nakagawa *et al*. (2007). Observation of GUS staining was conducted with an Olympus SZX10 microscope (Olympus Co. Ltd, Tokyo, Japan).

### Subcellular localization and proteins interaction analyses

The cDNA sequence of *PdeMIXTA04* (without stop codons) was cloned and recombined into the pBI121 vector to build the pBI121-PdeMIXTA04-GFP construct. Transient expression of the construct was conducted with four-week-old leaves of *Nicotiana benthamiana* mediated by *A. tumefaciens* GV3101. The transformed plants were cultured in dark for 24 h, following 24 h under light. Then, GFP fluorescence signal was examined by using a Zeiss LSM 710 confocal microscope (Zeiss Co. Ltd, Oberkochen, Germany).

To screen the proteins interacted with PdeMIXTA04, we built a cDNA library with carpel collected at −1, 1, 3, 5, 8, 15 DAP. The cDNAs were ligated into the pGADT7-Rec vector and transformed into yeast strain AH109. The bait vector of PdeMIXTA04 was built by recombining the *PdeMIXTA04* into the pGBKT7 vector fussed with GAL4-DNA binding domain. With the established bait vector and cDNA library, the interacting proteins were screened by the yeast two-hybrid (Y2H) following description by Tian *et al*. (2020). In this process, the pGBKT7-p53/pGADT7-T was used as the positive control and the pGBKT7-Lam/pGADT7-T was used as the negative control. Subcellular localization of the interacted proteins was performed as described above.

The screened interaction proteins were further verified by the Y2H, biomolecular fluorescence complementation (BiFC), dual-luciferase (dual-LUC), and pull-down experiments. In Y2H experiment, we used the Matchmaker GAL4 Two-Hybrid System and the experiment was performed following method described by Yu *et al*. (2010). For BiFC assay, pCAMBIA1300-35S-YFPc155 and pCAMBIA1300-35S-nYFP173 plasmids were used. The cDNAs (without stop codons) of *PdeMIXTA04* gene was fused to the N-terminus or C-terminus of the yellow fluorescent protein (nYFP or cYFP) to generate the plasmid constructs. Transient transformation of the established constructs was mediated by *A. tumefaciens* GV3101 with leaves from a four-week-old tobacco plant following descriptions by Xie *et al*. (2021). After 48 h, the YFP fluorescence signal was recorded using a Zeiss LSM 710 confocal microscope (Zeiss Co. Ltd, Oberkochen, Germany). For dual-LUC test, cDNAs (without stop codons) of the *PdeMIXTA04* genes were separately inserted into the N-terminus luciferase complementation image (nLUC) vector and the cLUC vector by using the Gateway method as described by Yu *et al*. (2020). Transient transformation of the established constructs was mediated by *A. tumefaciens* strain GV3101 with leaves from a four-week-old tobacco plant based on Yu *et al*. (2020). After 3 days, 10 μM luciferase substrate D-Luciferin (IL0230, Solarbio, USA) was sprayed on the surface of transformed leaves which were then incubated in the dark for 10 min. The luciferase bioluminescence images were captured by the dual-luciferase reporter assay system (GloMax 20/20 Luminometer; Promega, USA). For Y2H, BiFC, and dual-LUC assays, all experiments were repeated with three independent biological replicates. For pull-down examination, full-length cDNAs of the investigated genes were separately cloned into PGEX-6P-1 or pET-28a plasmids. Subsequently, the established constructs were transformed into *E. coli* to express glutathione S-transferase (GST) or Histidine (His)-labeled proteins were detected following description by Tian *et al*. (2020).

## Results

### Poplar catkin development

SEM examination revealed protrusions on epidermal cells of poplar placenta (placenta defined as the location where ovules are formed and which function to supply nutrients during the development and maturity of seeds (Griffiths *et al*., 2006)) at 1 DAP (**Fig. 1a**), while no protrusions were observed on samples collected at −1 DAP. It is noteworthy that the surface of every placental epidermal cell protruded at 1 DAP (**Fig. 1a**). It is also noteworthy that no protrusions were observed on the ovular epidermal cells (**Fig. 1a**). These observations suggest that female flower development on poplar catkins differs from the formation of cotton fibers in its origin and development. According to Haigle *et al*. (2012), cotton fibers are formed by protrusions of the ovular epidermal cells and partial ovular epidermal cells protrude to form cotton fibers. At 3 DAP, the protrusions in poplar elongated significantly and actively (**Fig. 1b**). At this stage, no apparent contraction of placenta was observed. At 5 DAP, the protrusions developed into long fibers and the hypertrophic placentas disappears completely, forming annular fiber appendages at the base of the embryo/seed which developed from the ovule (**Fig. 1c**).

**Fig. 1.**
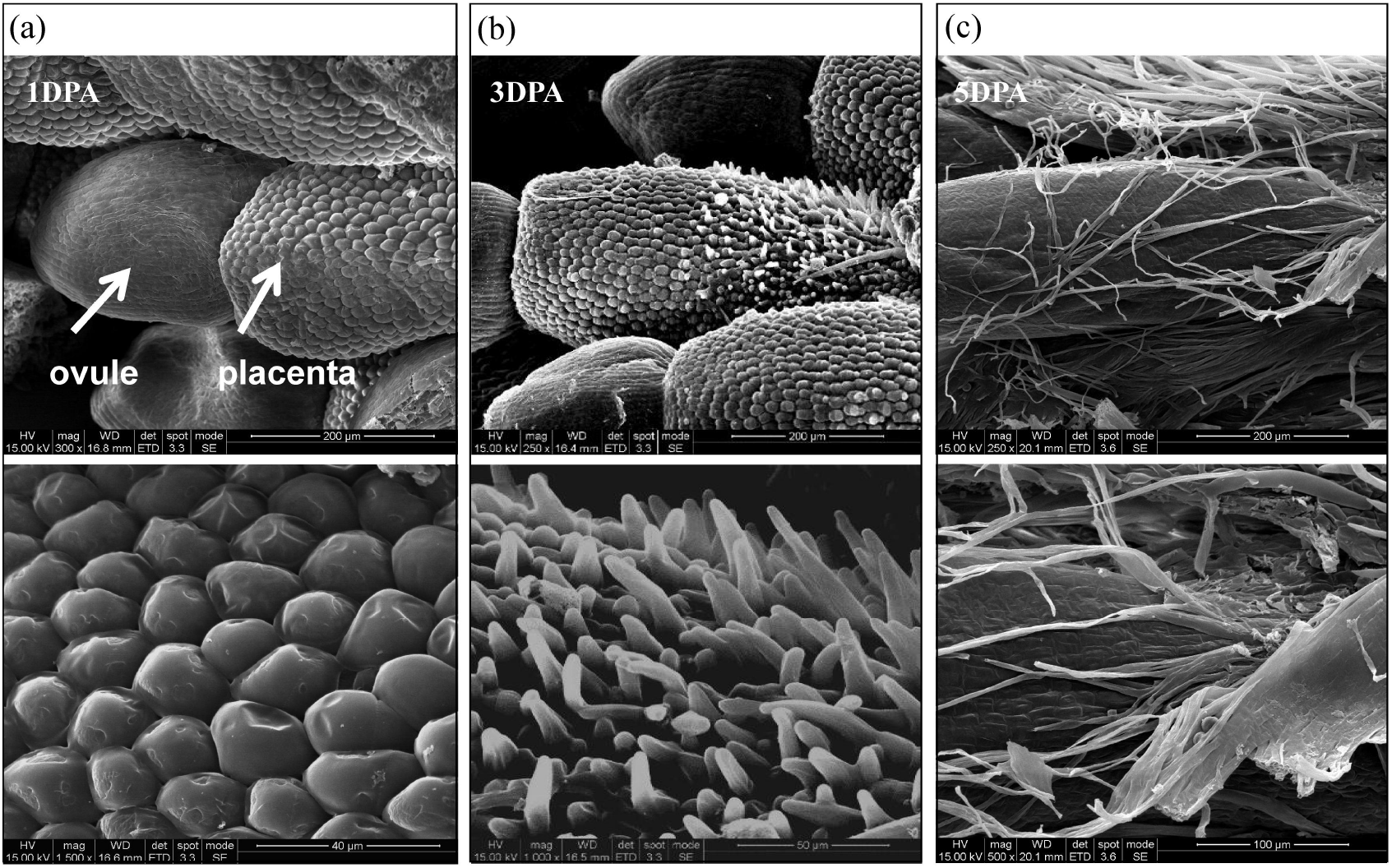
Poplar catkins development after pollination. **(a)** Protrusions on placental epidermal cells at 1 days after pollination (DAP), no protrusions were observed on ovular cells; **(b)** protrusions elongation at 3 DAP; **(c)** protrusions elongation at 5 DAP, placenta disappears at this stage. The upper and lower panels for **(a)**, **(b)**, and **(c)** represent the same image at alternate magnifications. Scale in the upper panel is 200 μm; scale in the lower panel is 40 μm.

To summarize SEM observations, female flower development occurs rapidly on the catkin immediately after pollination. Within 5 DAP, protrusion elongation of the placental epidermal cells leads to complete disappearance of the poplar placenta. During this process, cytokinesis was not observed and all fibers developed from a single cell. Contrary to the cotton fibers, flocculent poplar catkins developed from epidermal cells (ovule for cotton vs. placenta for poplar). Moreover, all placental epidermal cells develop protrusions that go on to form poplar catkin fibers. In contrast, in cotton, only partial ovular epidermal cells protruded to form cotton fibers (Stewart, 1975).

### Analysis of the *PdeMIXTA* genes

Based on homology to the conserved domains of MIXTA protein, eight *MIXTA* homologous genes, namely *PdeMIXTA01, PdeMIXTA02, PdeMIXTA03, PdeMIXTA04, PdeMIXTA05, PdeMIXTA06, PdeMIXTA07*, and *PdeMIXTA08*, were identified in the genome of *P. deltoides*. Subsequently, we cloned and resequenced these genes (GenBank accession numbers are OP476501, OP476502, OP476503, OP476504, OP476505, OP476506, OP476507, OP476508 https://www.ncbi.nlm.nih.gov/genbank/update.html). These *PdeMIXTA* genes all contained the conserved R2R3 domain and the AQWESAR**AE*RL*RES motif (**Fig. 2b**). Sequence alignment showed that *PdeMIXTA02* and *PdeMIXTA03* shared the highest identity of 95.01% among the eight homologs. Plotting the physical locations of *PdeMIXTA* genes (**Fig. 2a**) revealed that physical length between *PdeMIXTA02* and *PdeMIXTA03* was about 29 kb, indicating they were tandem duplicated genes, while all the other genes were positioned on alternate chromosomes. In the 12 species included for phylogenetic tree construction, a total of 75 *MIXTA* genes were identified. The established phylogenetic tree parsed these genes into clade I, clade II, and clade III (**Fig. 2c**). Among them, genes in clade III were the most well studied, with some confirmed to play a critically role in regulating the trichome development. For instance, the *MgML8* gene is reported to be associate with density of leaf trichomes in *Mimulus guttatus* (Scoville *et al*., 2011) and *AmMYBML1* is known to regulate trichomes and conical cell formation in *A. majus* (Lau *et al*., 2015). In wild-type *G. hirsute*, 10 *GhMML* genes were highly expressed during the initiation of cotton fiber development. When examined in the fiberless mutant of *Gossypium hirsute*, low expression was detected for six *(GhMML3, GhMML4, GhMML7, GhMML8, GhMML9*, and *GhMML10)* of the 10 expressed genes (Zhang *et al*., 2015). These six genes were found in clade III (**Fig. 2c**). Among which, *GhMML3* was identified as the key gene triggering development of “fuzz” fibers (Walford *et al*., 2011) and *GhMML4* as the key genes triggering development of “lint” fibers (Wu *et al*., 2018). For the *MIXTA* genes in *P. deltoides*, three of them, *PdeMIXTA02, PdeMIXTA03*, and *PdeMIXTA04*, were clustered in this subclade (**Fig. 2c**).

**Fig. 2.**
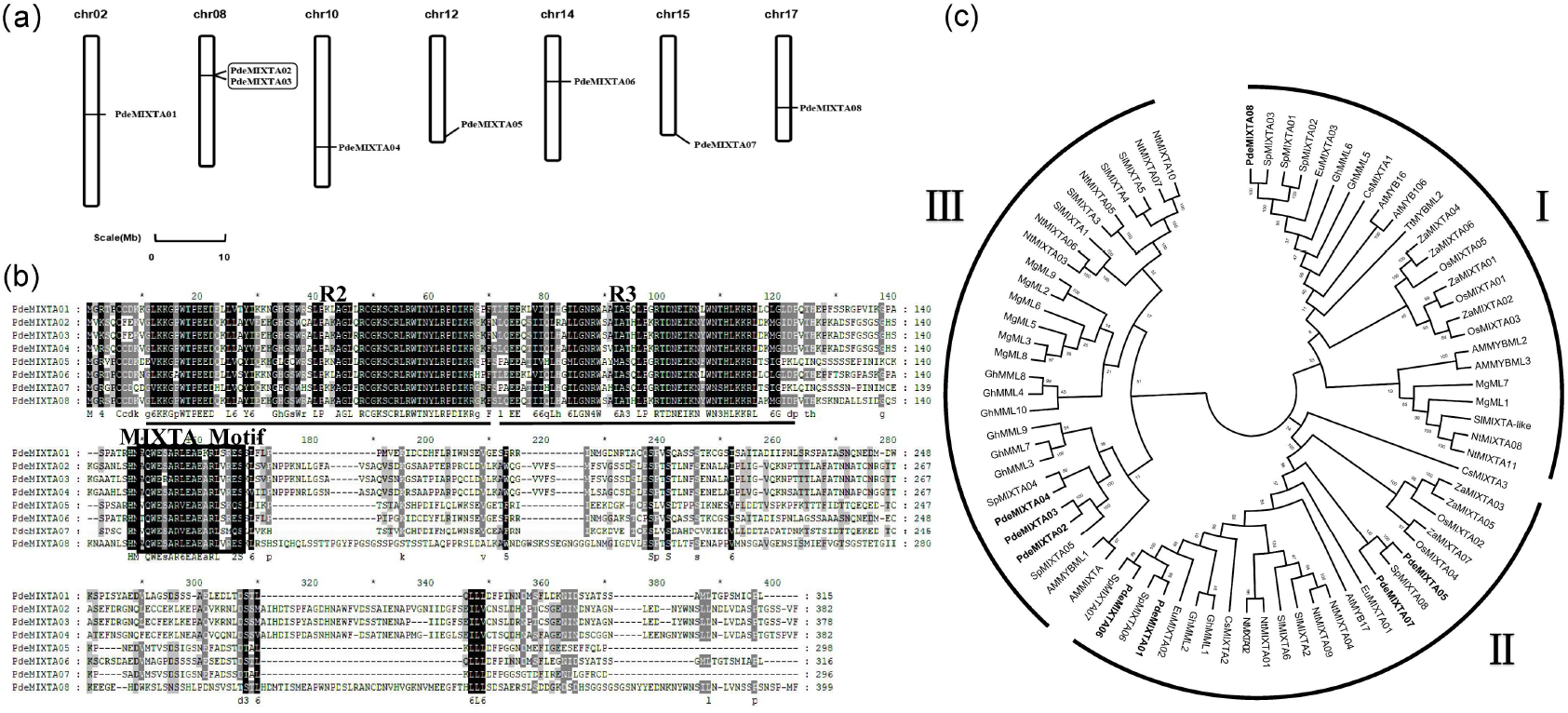
Chromosomal localization, sequence characteristics and phylogenetic analysis of the *PdeMIXTA* genes. **(a)** Chromosomal localization of the *PdeMIXTA* genes. The scale bar is shown at the bottom. **(b)** Sequence characteristics of the *PdeMIXTA* genes. **(c)** Phylogenetic analysis of *MIXTA* genes from 12 species. The *PdeMIXTA* genes were showed in bold.

### Comparative expression of the *PdeMIXTA* genes

With the carpel tissues of the female poplar flowers collected at −1, 1, 3, 5, 8, and 15 DAP, we analyzed the comparative dynamic expression of the *PdeMIXTA* genes from young leaves, vegetative buds, and bark tissues to test the tissue-specific expression of the *PdeMIXTA* genes (**Fig. 3a**). Based on the SEM observation, the underlying key genes were expected to be highly expressed in carpel of female poplar flower at the initiation stage of poplar catkins development. The expression of *PdeMIXTA02*, *PdeMIXTA03*, and *PdeMIXTA04* were consistent with this expectation, i.e., their expression was carpel-specific (**Fig. 3b**). As noted above, all the three genes are in clade III. By contrast, the expression of *PdeMIXTA01* (in clade I), *PdeMIXTA05* (in clade II), *PdeMIXTA06* (in clade II), *PdeMIXTA07* (in clade II), and *PdeMIXTA08* (in clade II) did not fit for the expectation of SEM observation and displayed relatively high expression in tissues other than the carpel (**Fig. 2c, 3b**).

**Fig. 3.**
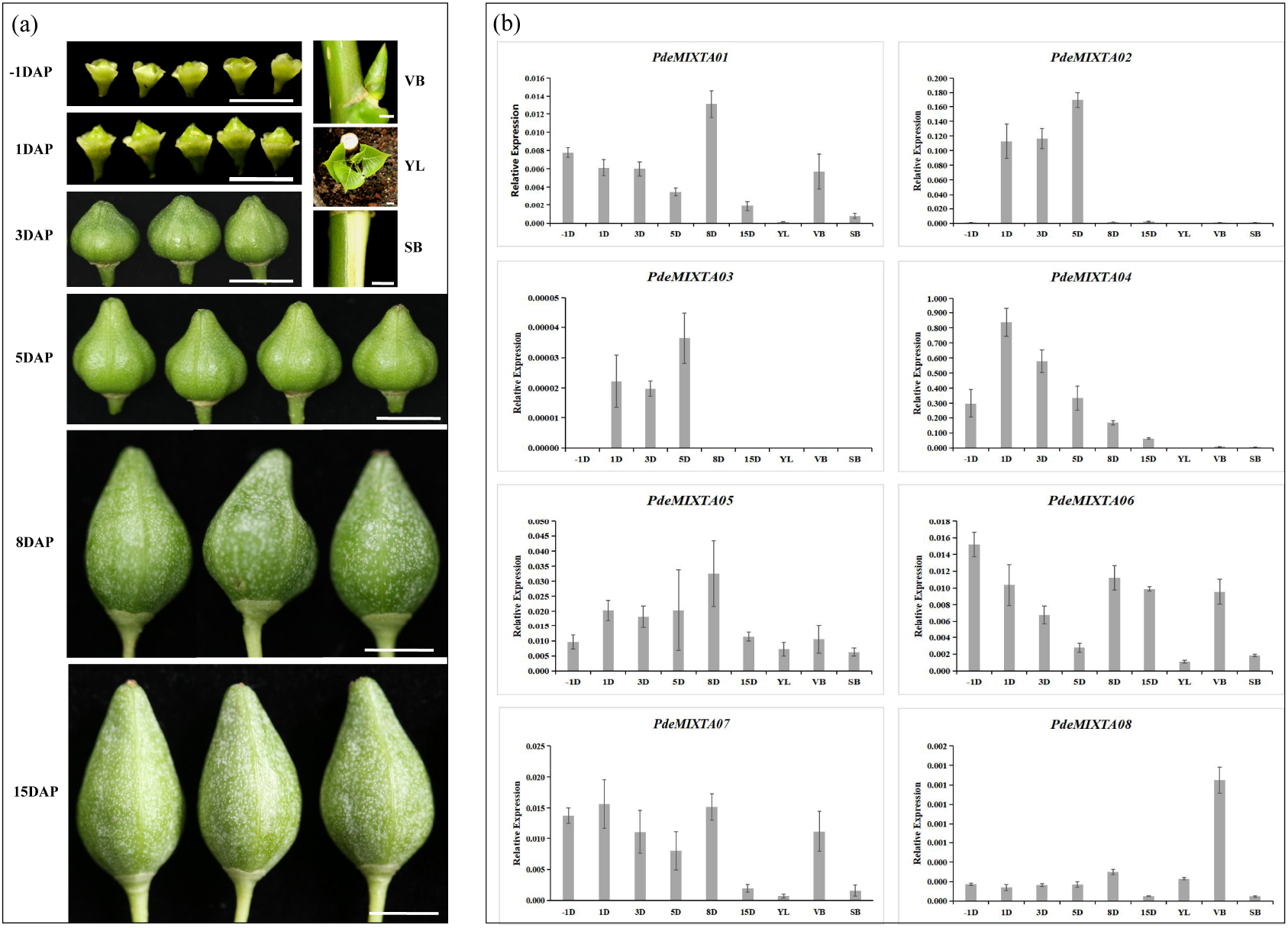
Development of carpel of the female poplar flowers and the differential expression of the *PdeMIXTA* genes. **(a)** Carpel morphology of female poplar flowers at −1, 1, 3, 5, 8, and 15 DAP, and the young leaves (YL), vegetative buds (VB), and stem bark (SB) used for qRT-PCR assay. **(b)** The relative expression level of *PdeMIXTA* genes determined by qRT-PCR. The vertical line on top of each column is the standard error bar (n=3).

Among the three carpel-specific genes, the tandem duplicated genes of *PdeMIXTA02* and *PdeMIXTA03* exhibited a similar differential expression pattern. Compared with *PdeMIXTA04*, *PdeMIXTA02* and *PdeMIXTA03* were expressed at a very low level after pollination, where expression levels of *PdeMIXTA04* was about seven times higher than that of *PdeMIXTA02*, and nearly 40,000 times higher than that of *PdeMIXTA03* at 1 DAP (**Fig. 3b**). Therefore, we hypothesized that *PdeMIXTA04* (the *Populus trichocarpa* homolog Potri.010G165700.1) plays a critical role in regulating the poplar catkins development. However, we could not exclude *PdeMIXTA02* and *PdeMIXTA03* merely based on their expression levels, and thus we included all the three carpel-specific genes from clade III as candidates for functional analysis in the subsequent transgenic studies.

### Functional analysis of the carpel-specific *PdeMIXTA* genes

*Arabidopsis* has been used to study trichomes developed from single-cell protrusions of the epidermis (Wang *et al*., 2019) and thus we tested *PdeMIXTA02, PdeMIXTA03* and *PdeMIXTA04* over-expression in *Arabidopsis*. For each gene, we analyzed 50 transgenic plants (five transgenic lines, with 10 plants for each line). The results showed that over-expression of *PdeMIXTA02* and *PdeMIXTA03* did not affect the development of leaf trichomes (**Fig. 4a**). By contrast, over-expression of *PdeMIXTA04* had significant effect on leaf trichomes. Statistically, trichomes density of the *35S:PdeMIXTA04* transgenic lines was 2.1 times higher than that of the wild-type plants (p-value<0.001) (**Fig. 4a**). Typically, leaf trichomes of wild-type *Arabidopsis* contain a large, polarized cell with two branch points and a large nucleus at the lower branch point, culminating in a trichome with three branches (Ishida *et al*., 2008). Based on SEM examination, the *35S:PdeMIXTA04* transgenic lines frequently displayed four or more trichome branches (**Fig. 4b**).

**Fig. 4.**
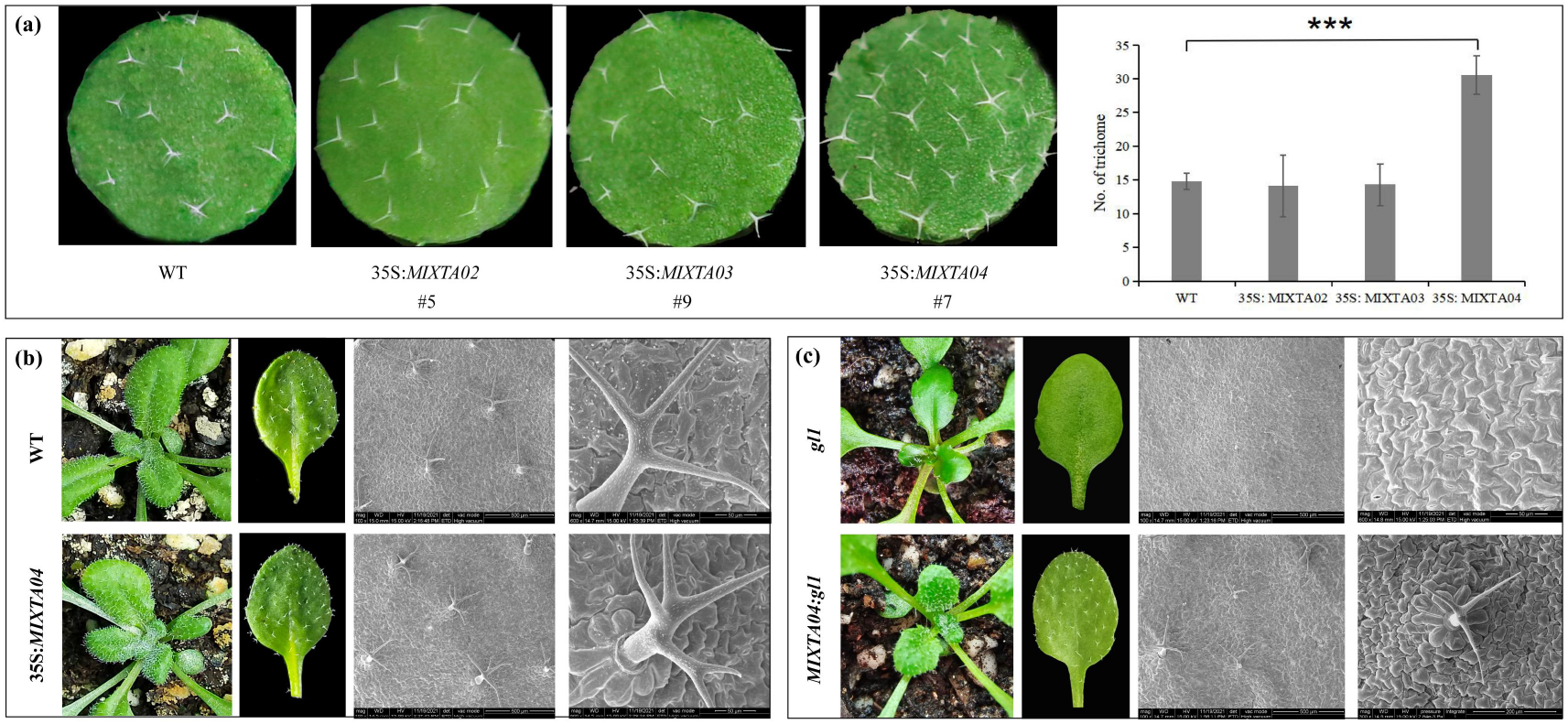
Ectopic expression of the three carpel-specific *PdeMIXTA* genes in *Arabidopsis*. **(a)** The rosette leaf trichome density of the wild-type plant, the *35S:PdeMIXTA02, 35S:PdeMIXTA03* and *35S:PdeMIXTA04* transgenic plants. The last panel at right showed the statistics of trichome numbers based on 50 transgenic plants. In this panel, the vertical bar on top of each column is the standard error bar. **(b)** Besides an increase in trichome density, overexpression of *PdeMIXTA04* also increased the trichome branches. **(c)** Ectopic expression of *PdeMIXTA04* recovered the trichome deficiency of *gl1* mutant. In **(b)** and **(c)**, the last panels at the most right showed the enlarged SEM scopes.

To further explore the functional nature of *PdeMIXTA04*, we transferred *PdeMIXTA04* into the trichome deficient mutants (*gl1*) of *Arabidopsis* (the upper panels of **Fig. 4c**) to see if *PdeMIXTA04* could complement the trichomeless phenotype of *gl1*. In sixteen hygromycin-resistant T1 transgenic lines, trichome development on rosette leaves was restored entirely (the lower panels of **Fig. 4c**), indicating ectopic expression of *PdeMIXTA04* could recover the trichome deficiency of *gl1* mutant. Thus, the transgenic studies have provided experimental evidence that, among *PdeMIXTA02, PdeMIXTA03* and *PdeMIXTA04, PdeMIXTA04* is the key gene regulating differentiation of epidermal cells.

### *PdeMIXTA04* specifically expressed in the poplar placental epidermal cells

SEM observations detected the initiation of protrusions on top of the epidermal cells of poplar placenta at 1 DAP, which was also observable under microscope with the longitudinal sections of poplar female flower collected at 1 DAP (**Fig. 5a**). With slices prepared at this stage, we conducted *in situ* hybridization to corroborate the specific expression of *PdeMIXTA04* in epidermal cells of poplar placentas. Hybridized with the antisense and the sense probes, which were designed according to the specific sequence of the 3′ end of *PdeMIXTA04*, a clear hybridization signal occurred only in the protrusions of the placental epidermal cells with the antisense probe (**Fig. 5b**), indicating that *PdeMIXTA04* is explicitly expressed in the epidermal cells of poplar placentas.

**Fig. 5.**
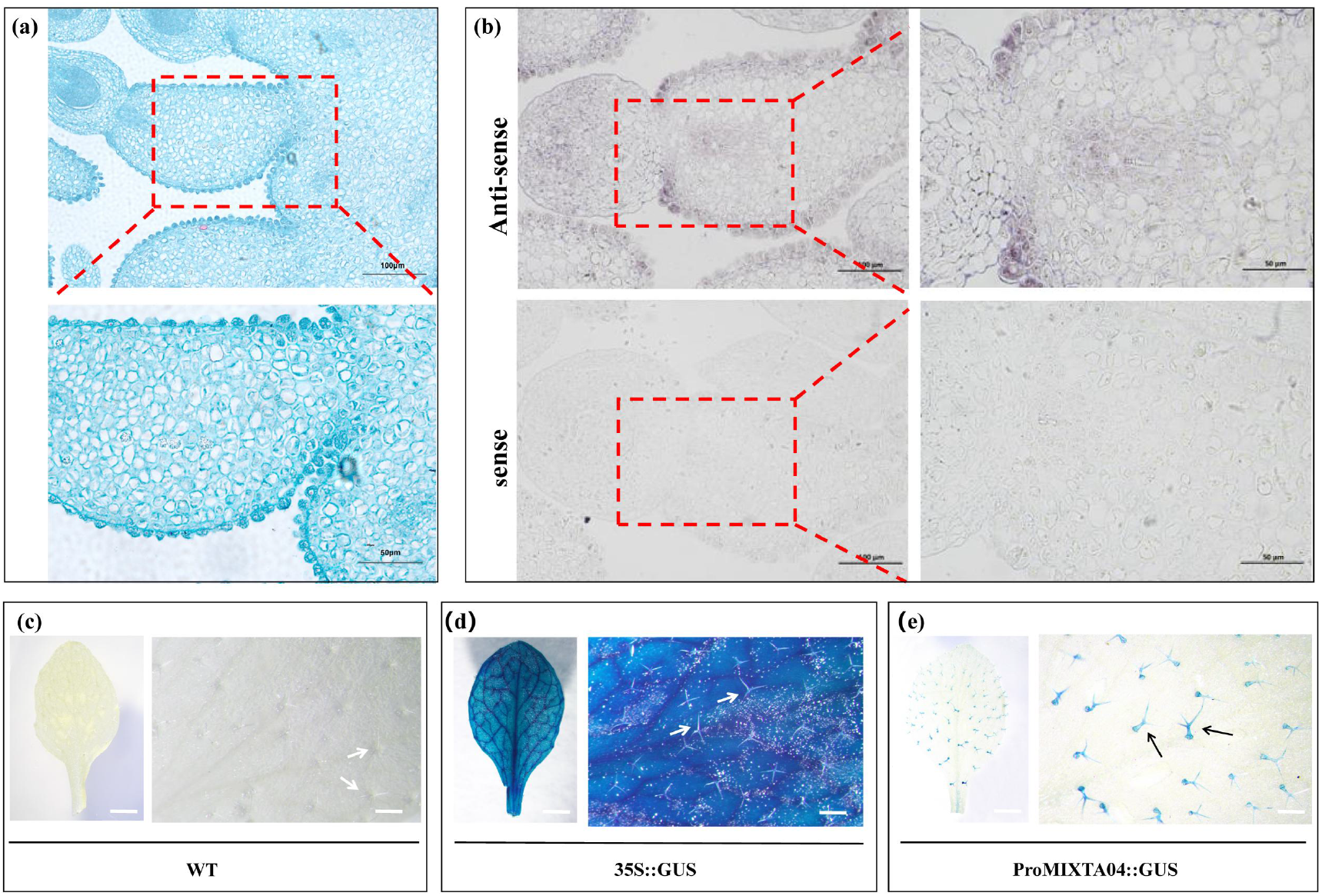
*PdeMIXTA04* specifically expressed in the epidermal cells of poplar placentas and its promoter mediated trichome-specific expression. **(a)** The longitudinal section of poplar female flower collected at 1 DAP. The scale bar of the upper panel is 100 μm, and that of the lower panel (enlarged scope) is 50 μm; **(b)** *in situ* hybridization with the antisense probe (the upper panels) and the sense probe (the lower panels) for *PdeMIXTA04* at 1 DAP. The scale bar of the left panels is 100 μm, and that of the right panels (enlarged scopes) is 50 μm; **(c)-(e)** showed the GUS staining results: **(c)** GUS expression in the wild-type plants (negative control). **(d)** GUS expression in transgenic plants of 1301-35S-GUS (positive control). **(e)** GUS expression in transgenic plants of 1301-ProMIXTA04. The scale bar of the left panels is 5 mm, and that of the right panels (enlarged scopes) is 0.5 mm.

To further define epidermal cell-specific expression of *PdeMIXTA04*, we cloned the promoter of *PdeMIXTA04* (GenBank accession No. OP493866) and constructed a 1301-ProMIXTA04-GUS vector to create a GUS report gene. The 1301-ProMIXTA04-GUS vector and 1301-35S-GUS vector were independently transformed into the wild-type *Arabidopsis*. Leaves of the two-week-old plants were used for GUS staining analysis. No GUS expression was detected in the wild-type (negative control) (**Fig. 5c**). In the 1301-35S-GUS transgenic plants (positive control), GUS expression covered the entire leaf (**Fig. 5d**). In contrast, GUS expression was observable only in the leaf trichomes of 1301-ProMIXTA04-GUS transgenic plants (**Fig. 5e**). Based on the above observations, GUS staining clearly demonstrated that promoter of *PdeMIXTA04* mediated trichome-specific expression.

### *PdeMIXTA04* function through the MBW complexes

Subcellular localization of PdeMIXTA04 via transient expression in *N. benthamiana* demonstrated that *PdeMIXTA04* localized to the nucleus (**Fig. S2**), suggesting *PdeMIXTA04* may function as a transcription factor alone or in a complex. Screened with pGBKT7-PdeMIXTA04 bait vector against the carpel-specific yeast library, we observed both the *bHLH (PdeMYC)* and *WD40 (PdeWD40)* genes interacted with *PdeMIXTA04*, which differs from the findings in cotton, where the *MIXTA*-like gene (*GhMML4*) was observed to intact only with *WD40* (*GhWDR*) and not *bHLH* (Tian *et al*., 2020). However, other *MIXTA* genes have been reported to function through interaction with *bHLH* and *WD40* genes in almost all investigated plants, *e.g*., *A. thaliana* (Zhao *et al*., 2008), *Actinidia chinensis* (Liu *et al*., 2021), and *Camellia sinensis* (Li *et al*., 2022). These reports and our results suggest that PdeMIXTA04 of poplar functions through the common MBW complexes retained in most plants.

We further conducted the Y2H, BiFC, dual-LUC, and pull-down experiments to confirm the interactions of PdeMIXTA04 with PdeMYC and PdeWD40. With PdeMIXTA04 and PdeMYC fused to the GAL4 DNA binding domain (BD) and PdeWD40 and PdeMYC fused to the activation domain (AD), the Y2H assay demonstrated that yeast transformed with both vectors could grow on medium without tryptophan, leucine, histidine, and adenine (**Fig. 6a**). Y2H results, presented in the 1^st^, 2^nd^, and 3^rd^ lines of **Fig. 6a**, confirmed the interactions between PdeMYC and PdeMIXTA04, between PdeWD40 and PdeMIXTA04, between PdeMYC and PdeWD40, respectively, indicating that these three proteins interact in a complex. In the BiFC assay, CDS sequences of *PdeMIXTA04* and *PdeWD40* were separately fused to the nYFP fragment and CDS sequences of *PdeMYC* and *PdeWD40* were separately fused to the cYFP fragment. Transient transformation of these vectors in tobacco leaf cells produced visible fluorescence signals (**Fig. 6b**). The BiFC assay results in the upper panel, the middle panel, and the bottom panel of **Fig. 6b** clearly showed the interactions between PdeMIXTA04 and PdeWD40, between PdeMIXTA04 and PdeWD40, between PdeWD40 and PdeMYC, respectively. Subcellular localization of PdeWD40 and PdeMYC detected both nuclear and membrane localization signals (**Fig. S2**), which confirmed that the observed BiFC signals were not false positive. In dual-LUC assay, CDS sequences of *PdeMIXTA04* and *PdeWD40* were separately fused to the nLUC and *PdeMYC* and *PdeWD40* were separately fused to the cLUC. Visible luciferase activity was evident in transient transformation of PdeMIXTA04-PdeMYC, PdeMIXTA04-PdeWD40, and PdeMYC-PdeWD40 vectors combination in tobacco leaf cells (**Fig. 6c**), which confirmed the pair-wise interactions of the three proteins. Finally, the protein interactions were further verified by the *in vitro* pull-down experiment (**Figure 6d**). Incubation of GST-PdeMIXTA04 with His-PdeMYC (the upper panel) showed that GST-PdeMIXTA04 could effectively pull down His-PdeMYC. By contrast, the negative control (GST resin) could not pull down the His-PdeMYC. These results indicated that PdeMIXTA04 physically interacted with PdeMYC *in vitro*. In the pull-down experiment, incubation of GST-PdeWD40 with His-PdeMYC, and incubation of GST-PdeWD40 with His-PdeMIXTA04 also confirmed the physically interactions between PdeWD40 and PdeMYC (the middle panel) and between PdeWD40 and PdeMIXTA04 (the bottom panel). In total, based on the above experiments, we conclude that there is a tripartite interaction among PdeMIXTA04, PdeMYC and PdeWD40, both *in vivo* and *in vitro*, which supports our hypothesis that PdeMIXTA04 of poplar functions through the MBW complexes.

**Fig. 6.**
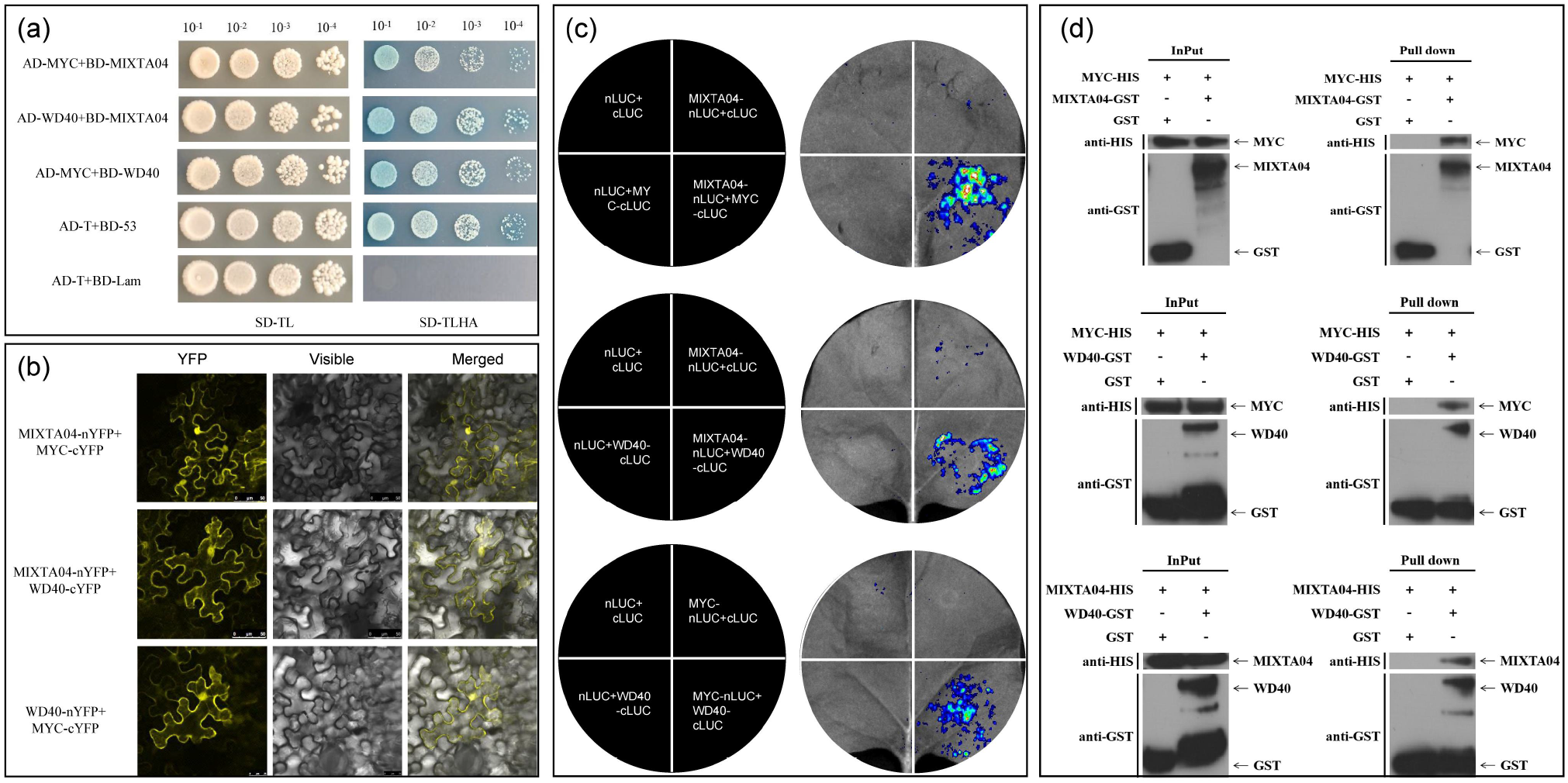
Experimental results confirmed the interactions among proteins coding by *PdeMIXTA04*, *PdeMYC*, and *PdeWD40* genes. **(a)** The Y2H assay confirmed the PdeMYC-PdeMIXTA04 (1^st^ line), PdeWD40-PdeMIXTA04 (2^nd^ line), and PdeMYC-PdeWD40 (3^rd^ line) interactions. The fourth line is the positive control and the fifth line is the negative control. AD represents the prey vector and BD represents the bait vector. The left panel showed the yeast cells in dilution gradients of 10^−1^, 10^−2^, 10^−3^, and 10^−4^ cultured on the SD/-Trp/-Leu (SD/-TL) medium. The right panel showed the yeast cells in dilution gradients of 10^−1^, 10^−2^, 10^−3^, and 10^−4^ cultured on the SD/-Trp/-Leu/-His/-Ade (SD/-TLHA) medium. **(b)** The BiFC assay confirmed the PdeMIXTA04-PdeMYC (the upper panel), PdeMIXTA04-PdeWD40 (the middle panel), and PdeWD40-PdeMYC (the bottom panel) interactions. Presence of yellow fluorescence indicated the corresponding proteins interacted with each other. YFP, yellow fluorescent marker; Visible, bright-field; Merged, merged view of YFP and Visible. **(c)** The dual-LUC assay confirmed the PdeMIXTA04-PdeMYC (the upper panel), PdeMIXTA04-PdeWD40 (the middle panel), and PdeMYC-PdeWD40 (the bottom panel) interactions. The left panel showed the construct combinations, and the right panel showed the fluorescent signals on the corresponding parts of the injected leaves. For different construct combinations, those containing interacted proteins could induce fluorescence (the lower left portion of the leaves). **(d)** The pull-down assay confirmed the interaction between PdeMYC and PdeMIXTA04 (the upper panel). In which, the “Input” showed the gel electrophoresis of the corresponding input proteins (control). Confirmation of interaction between PdeMYC and PdeWD40 was shown in the middle panel, and that between PdeMIXTA04 and PdeWD40 was shown in the bottom panel.

As noted above in cotton, *GhMML4* interacts with *GhWDR* (*WD40*) to regulate cotton fiber development (Tian *et al*., 2020). Phylogenetic analysis showed that *PdeWD40* and *GhWDR* were clustered into the same *WD40* subclade (**Fig. S3a**). Moreover, structural analysis showed that GhWDR and PdeWD40 possessed the same number of WD40 repeat domains (**Fig. S3b**), suggesting *PdeWD40* and *GhWDR* might be functionally conserved. In *Arabidopsis, AtTTG1* is a *WD40* gene that interacts MYB gene (*GL1*) to modulate leaf trichome development (Zhao *et al*., 2008). Contrary to the *PdeWD40* and *GhWDR* genes, *AtTTG1* was clustered into a different subclade (**Fig. S3a**) and has fewer WD40 repeat domains, implying *AtTTG1* might be functionally distinct from *PdeWD40* and *GhWDR*. As for bHLH in the MBW complex, no *bHLH* gene was observed to interact with *GhMML4* or *GhWDR* in cotton (Tian *et al*., 2020). However, in *Arabidopsis, AtGL3*, a *bHLH* gene, interacts with *GL1* and *AtTTG1* (Esch *et al*., 2003). Structure analysis showed that *AtGL3* and *PdeMYC* shared the same conserved domains (**Fig. S3b**), suggesting that the *bHLH* gene in poplar and *Arabidopsis* might be functionally conserved, supporting our hypothesis that PdeMIXTA04, PdeMYC and PdeWD40 form a functional complex.

## Discussion

### Placental epidermal cells protrude collectively to form catkins immediately after pollination

The placenta in fruits of most plants eventually desiccate and shrink as the fruit matures. In addition to giving nutrients for seed development, the placenta of some plants develop further and participate in the formation of fleshy tissue, such as the fruit of tomato and kiwi (Zhang *et al*., 2019b; Guo *et al*., 2013). The placenta of other plants develop into thick, fleshy aril that encase the developing seeds, *e.g*., in litchi, longan, durian, and others (Zhang, 2001). The entire ovary of *Populus* species consists of an ovule and a placenta attached at the base, and the placental epidermal cells protrude to ultimately form unisexual apetalous flowers arranged in a catkin (Ye *et al*., 2014). Thus far, the detail of the development of poplar catkins has been poorly characterized. In this study, our SEM observation provided insight into the development of poplar catkins. These images clearly showed that trichomes of the female flower of poplar occur as a collection of single-cell protrusions of the placental epidermal cells. Female catkin in poplar development was rapid, initiating immediately after pollination with all placental epidermal cells display protrusion that ultimately form unisexual apetalous flowers arranged in a catkin. The placental tissues disappears at 5 DAP leaving elongated fibers. Female catkins in poplar are superficially similar as the cotton fibers, where both function as seed hair to assist wind dispersal. Thus, female catkins in poplar and cotton fibers have been considered to be generated through the same molecular apparatus (Stewart, 1975; Zhang *et al*., 2015). However, cotton fibers originate from the ovular epidermal cells and only partially protrude to form cotton fibers (Wu *et al*., 2018). In contrast, in poplar, the placental tissue generated a distinctive annular fiber appendage at the radical end of carpel, with the placenta disappearing by the end. These observations suggest that there is a difference in the regulation of seeds hair development between the two species.

### *MIXTA* genes regulate plant epidermal cell differentiation

Plant *MIXTA* genes encode transcription factors and these genes belong to the SBG9 subgroup of the *R2R3 MYB* gene family (Brockington *et al*., 2013). Besides the R2R3 domain, the *MIXTA* genes are characterized with an additional conserved motif AQWESAR**AE*RL*RES, a 10-22 amino acids series (Grotewold, 2006). It has been reported that *MIXTA* genes play critical roles in regulating differentiation of epidermal cells of different plant tissues, such as root hairs, conical cells, and leaf trichomes (Shi *et al*., 2018). In some plant lineages *MIXTA* genes are involved in the regulation of epidermal cells. For instance, in *A. annua AaMIXTA1* regulates the formation of glandular secretory trichomes (Espley *et al*., 2007) and in *G. hirsuta GhMML3* and *GhMML4* triggers the development of fuzz and lint fibers, respectively (Walford *et al*., 2011; Wu *et al*., 2018) and in *A. thaliana AtMYB106* modulated leaf trichome development (Zhao *et al*., 2008).

We identified eight *MIXTA* homologous genes in *P. deltoides* genome, each containing the characteristic AQWESAR**AE*RL*RES motif. Phylogenetic analysis clustered these eight poplar *MIXTA* genes into three subclades. The *PdeMIXTA08* were clustered in the same clade as *AtMYB106*, suggesting that *PdeMIXTA08* might be a *MIXTA* gene involved in poplar leaf trichome development. It was noteworthy that *PdeMIXTA02*, *PdeMIXTA03*, and *PdeMIXTA04* were clustered in the same clade as *GhMML3* and *GhMML4*, suggesting the three genes might involve in the formation of poplar seed hairs. Based on these relationships and through a series of experiments, we confirmed that *PdeMIXTA04* was a lineage-specific *MIXTA* gene in poplar that triggered early development and subsequent formation of female catkins in poplar. *Salix* (willow) is the sister genera of *Populus*, also produces catkins. Within the phylogenetic tree, the willow *SpMIXTA04* gene was the most closely related to *PdeMIXTA04*. It is likely that the formation of willow and poplar catkins is likely triggered by the same molecular apparatus.

### Poplar retains both partner proteins in the MBW complex

In plants, previous studies have discovered a variety of genes either positively or negatively regulate patterning of surface hairs of different plant tissues. A regulatory system composed of multiple transcription factors appears crucial to trichome differentiation and development (Wang *et al*., 2019). Regulation of leaf trichomes development by the MYB-bHLH-WD40 (MBW) was first described in *Arabidopsis* (Zhao *et al*., 2008), thereafter, this complex was reported to control the spatial positioning of trichomes and their density on leaves in various other plants (Zhang *et al*., 2019a, Li *et al*., 2022, Shi *et al*., 2018). For example, in the tea plant, CsMYB1 was found to interact with CsGL3 and CsWD40, forming the MBW transcriptional complex to activate the trichome development associated genes, *CsGL2* and *CsCPC*, resulting in formation of trichomes on leaves (Li *et al*., 2022). In our study, we characterized the partner proteins, PdeMYC and PdeWD40, in the MBW complex of poplar using PdeMIXTA04 as bait and experimentally confirmed the mutual interactions among the three proteins. The MBW complex typically consisted of three proteins encoded by a *MYB*, a *bHLH* and a *WD40* gene. To date, the absence of protein encoded by the *bHLH* gene in the MBW complex was only reported in *G. hirsutum*. In our study, we found that the development of female flowers arranged in catkins in poplar was triggered by the *PdeMIXTA04* gene, which is closely related to the *GhMML3* and *GhMML4* genes in cotton and more divergent from the *AtMYB106* gene in *Arabidopsis*. In terms of the MBW complex, unlike cotton, poplar retained both partner proteins. The comprehensive work carried out in this study have added to the growing body of evidence that the *MIXTA* genes regulate plants epidermal cell differentiation and provided unique perspectives for better understanding the molecular mechanism underlying plant seed hair development.

## Consent for publication

All authors have read and approved the manuscript

## Supplementary files

**Table S1**. Primers used for cloning the *PdeMIXTA* genes, gene-specific primers used for qRT-PCR assay, and probes used *in situ* hybridization.

**Fig S1**. Circular map of the 1301-ProMIXTA04-GUS construct.

**Fig S2**. Subcellular localization of *PdeMIXTA04*, *PdeWD40* and *PdeMYC*. The scale bar is 25 μm.

**Fig S3**. Phylogenetic analysis and structural comparison of the *WD40* and *bHLH* genes involving in MBW complex. **(a)** Phylogenetic analysis of the *WD40* involving in MBW complex in *Populus deltoides*, and *Gossypium hirsute*. Tree construction was performed following the same computational pipeline as that of Figure 2c. **(b)** Structural comparison of the *WD40* and *bHLH* genes involving in MBW complex among the above three species. The grey boxes represent the characteristic domains of the corresponding genes. For the *WD40* genes, *PdeWD40* in *P. deltoides* and *GhWDR* in *G. hirsute* both contain six WD40 repetitive domains, while *AtTTG1* in *A. thaliana* only possesses four WD40 repetitive domains. For the *bHLH* genes, *PdeMYC* in *P. deltoides* and *AtGL3* in *A. thaliana* share the same characteristic domains. No *bHLH* gene was detected to interact with *WD40* and *MYB* in *G. hirsute*.

